# Alzheimer’s Disease Brain Organoids as a Source of Disease-Relevant Amyloid-Beta Oligomers

**DOI:** 10.64898/2026.03.09.710594

**Authors:** Eric Zanderigo, Manaal Fatima, Sandy Becker, Alison O’Neil

**Affiliations:** Department of Chemistry, Wesleyan University, Middletown, CT, USA; Department of Biology, Wesleyan University, Middletown, CT, USA; Department of Chemistry, Neuroscience and Behavior Program, Wesleyan University, Middletown, CT, USA

**Keywords:** Alzheimer’s Disease, amyloid beta, brain organoids

## Abstract

Amyloid plaques are a hallmark of Alzheimer’s disease (AD) progression; however, the early stages of plaque formation and the specific amyloid-beta (Aβ) species involved remain difficult to study. While post-mortem tissue provides insight into end-stage mature plaques, therapeutic development requires targeting the earliest Aβ oligomers to arrest plaque formation. Furthermore, inherently toxic soluble Aβ oligomers off-pathway from plaque formation are implicated as a driving force of AD pathology. It also remains unclear if the specific nature of key disease-relevant species can be accurately replicated in preparations of synthetic peptides.. To bridge this gap, we utilize brain organoids carrying AD mutations as a biologically authentic source for Aβ peptides and oligomers. We demonstrate that these mutations do not disrupt organoid development and that the resulting conditioned media contains Aβ oligomers with disease-relevant structures. Finally, we show that these oligomers can be concentrated and segregated via differential ultracentrifugation for further experimental applications.

## 1. Introduction

The majority of global dementia cases are caused by Alzheimer’s disease (AD). The disease is characterized in part by the presence of intracellular neurofibrillary tangles (NFTs) of hyperphosphorylated tau and extracellular plaques of amyloid beta peptide (Aβ), both of which exert a toxic effect on the brain leading to the progressive death of neurons and decline of cognitive function. Many AD studies rely on the analysis of postmortem tissue; however, these samples inherently reflect end-stage disease conditions and protein populations. To instead model AD *in vitro*, human induced pluripotent stem cell (iPSC)-derived neuronal cultures have become increasingly common. While there is a wide variety of AD iPSC based models, the majority of studies have used 2D neuronal cultures^1^. Importantly, AD pathology and the bioactivity of disease-associated proteins is inherently tied to the complex cellular niche in which they are generated, so to more accurately model early stage AD, there is a growing emphasis in the field on iPSC-derived 3D cerebrocortical organoids. These complex tissue models are characterized by their brain-like cellular diversity and cytoarchitecture and are therefore uniquely poised to capture the *in vivo* conditions of AD pathologies in the brain. The presence of diverse cell types and ordered cortical layers in organoids ensures greater physiological relevance of the bioactive forms of AD-associated proteins than other models^1–3^. While current organoid studies primarily focus on intratissue accumulations, secreted soluble Aβ species, particularly neurotoxic soluble oligomers of Aβ_42_, play a pivotal role in driving AD pathophysiology^1,4,5^. The secreted proteomes (“secretomes”) of brain organoids, however, are an underutilized resource and have not been fully assessed as a scalable source for disease-relevant neurotoxic Aβ oligomers (AβOs).

As the use of brain organoids to model AD grows, standard protocols need to be established to ensure repeatability and rigorous conclusions. Since there are several approaches to organoid preparation and their associated expression of phenotypes vary, it is critical to specifically define AD hallmarks such as the amyloid plaque burden and the extracellular soluble Aβ populations associated with cultures generated through distinct differentiation protocols. The present study aims to both further our understanding of the intra- and extra-tissual properties of established AD cerebrocortical organoid models, and also to explore the untapped potential of these cultures as a physiologically relevant scalable source for bioactive soluble AβOs secreted by the brain in the early stages of AD. Our findings indicate a differentiation stage-linked flux in Aβ expression using several metrics for quantifying amyloid load in organoid sections and conditioned media and highlights that amyloid-burden alone should not be used to define the diseased state.

We successfully derived cerebrocortical organoids from an engineered mutant AD iPSC line (DKI, NYSCF BN0002, APP Swe/Swe, PSEN1 M146V/M146V) with its isogenic control (WT, NYSCF 7889SA), and an AD patient line (UCSD; APP duplication) using two distinct protocols. We discovered that our WT line produced Aβ plaques *within* the organoid in similar amounts to our disease lines, reflecting a known feature of the non-demented human population: about 30% of cognitively normal adults have significant amyloid plaque buildup^6,7^. In contrast, however, we characterized all lines’ secreted Aβ profiles through analyses of conditioned media and found distinctions between the diseased and WT lines. In the conditioned media, ELISA analyses showed elevated Aβ_42_ and Aβ_40_ levels, and elevated Aβ_42_/Aβ_40_ ratios in diseased lines and, by using cushioned differential ultracentrifugation (CD-UC) methods, we detected the production of AD-associated AβOs in the DKI line and isolated them from the bulk media. These results highlight the importance of the soluble secreted component of organoid cultures in AD research and lay the groundwork for using ultracentrifugation-based methods to detect and isolate patient-specific pathogenic Aβ species from AD organoid conditioned media for further characterization.

## 2. Materials and Methods

### iPSC Maintenance

Induced pluripotent stem cells (iPSCs) were maintained under feeder-free conditions (Corning Matrigel hESC Qualified Matrix, cat. no. 354277) using mTeSR™ Plus medium (STEMCELL Technologies, cat. no. 100-0274) supplemented with 10% mTeSR™ Plus Supplement (STEMCELL Technologies, cat. no. 100-0275). Cultures were incubated at 37°C with 5% CO₂ in a humidified incubator. Medium was refreshed daily, and cells were passaged as clumps upon reaching approximately 80% confluency using 0.5 mM EDTA.

### Organoid Generation

Cortical organoids were generated following the protocol of Velasco et al^8^, with minor modifications. Once iPSCs reached about 80% confluency, they were dissociated into single cells and adjusted to a density of 1x10⁶ cells/mL. Approximately 5x10⁶ cells in 5 mL of media were seeded per well in a six-well plate and maintained on an orbital shaker in a humidified incubator at 37°C with 5% CO₂. Media changes and small-molecule additions were performed according to the original protocol, except that organoids were maintained in the same plates throughout differentiation rather than being transferred to spinner flasks.

Organoids were also generated using a modified version of the Whye et al. protocol^9^. iPSCs were seeded as described in the Whye method, with the following adjustments: both the neural differentiation medium and the neural induction basal medium contained 0.055 mM 2-mercaptoethanol, instead of the suggested 0.1 mM. In addition, the small-molecule inhibitor DAPT was omitted from the Stage 4 cortical differentiation termination medium as it is a beta secretase inhibitor and could interfere with the production of Aβ species. At day 35, the resulting organoids were transferred to cortical neuron maturation medium and maintained at 37°C and 5% CO₂ with continuous shaking at 120 rpm until further processing.

### Lysis and Protein Extraction

At the designated time of harvesting, approximately 500 µl of organoid suspension was taken out from the rocking dish. Organoids were then washed three times with phosphate-buffered saline (PBS) and resuspended in RIPA buffer (Thermo Scientific, cat. no. 89900) supplemented with HALT protease and phosphatase inhibitors (Thermo Scientific, cat. no. 1861280) and Benzonase nuclease (Sigma-Aldrich, CAS 9025-65-4). Samples were mechanically disrupted using a pestle and incubated on ice for 30 minutes. Lysates were subjected to three freeze-thaw cycles using liquid nitrogen and a 37°C water bath, followed by centrifugation at 16,000 × g for 10 min at 4°C. The supernatant was collected and stored at -80°C as total soluble protein lysate.

### Organoid cryosectioning

Organoids were collected at the designated developmental day, washed three times with PBS, and fixed in 4% paraformaldehyde (PFA; Electron Microscopy Sciences, cat. no. 15710) for at least 4 hours at room temperature or overnight at 4°C. Fixed organoids were transferred to 30% sucrose in PBS and incubated overnight at 4°C until fully equilibrated, as indicated by sinking to the bottom of the tube. Organoids were then embedded in 10% gelatin in PBS and incubated at 37°C for 45 to 60 minutes. Custom molds were prepared using aluminum foil wrapped around 1.5 mL microtubes. The gelatin embedded organoids were transferred into the molds, allowed to settle for 1 minute, and flash-frozen in liquid nitrogen. Blocks were labeled and stored at -80°C until cryosectioning.

Frozen organoids were mounted onto cryostat chucks using OCT Compound (Fisher Healthcare, cat. no. 0004585/Rev B). Samples were equilibrated on dry ice for 10 to 15 minutes and then placed inside the cryostat chamber to acclimatize to the set temperature (-25°C chamber, -30°C chuck). After 30 minutes of stabilization, 15 µm sections were obtained using a cryostat (Leica CM series) and collected on Superfrost glass slides (Globe Scientific, cat. no. 1358W). Every ninth section was mounted together on a single slide, resulting in 4 to 6 sections per slide. Prepared slides were stored at -20°C until stained.

### Immuno-labeling and Chemical Staining

Slides were brought to room temperature and washed with PBS at 37°C for 10 minutes. Sections were blocked for 1 hour in a solution containing 5% bovine serum albumin (BSA; Fisher BioReagents, cat. no. BP9703-100) and 5% horse serum in PBS with 0.1% Triton X-100. Primary antibodies were diluted in primary dilution buffer consisting of 5% BSA and 0.05% sodium azide in PBST (PBS with 0.1% Tween® 20) and applied overnight in a humidified chamber at 4 °C. Antibodies used were TBR1 (Abcam, cat. no. Ab183032), CTIP2 (Abcam, cat. no. ab18465), SATB2 (Abcam, cat. no. Ab51502), MAP2 (Invitrogen, cat. no. PA1-16751), Aβ 6E10 (Biolegend, cat no. 803002). After incubation, slides were washed three times in PBST for 5 minutes each and incubated with appropriate fluorophore-conjugated secondary antibodies diluted in blocking buffer for 2 hours at room temperature. Following three 30-minute PBST washes, slides were air-dried and cover slipped with ProLong™ Gold antifade reagent (Invitrogen, cat. no. P36934). AmyloGlo (Biosensis, cat. no. TR-400-AG) and Proteostat staining (Enzo, cat. no. ENZ-51035-0025) of organoid sections was carried out as described by the manufacturer.

### Confocal Imaging

Images were acquired using a Leica SP8 confocal microscope. Scans were captured at 1024 x 1024 pixels using either a 20x dry objective or a 63x oil-immersion objective. Z-stacks were collected with a 1 µm step size across a total depth of 15 µm. Maximum intensity projections were generated using Leica LAS X software, with a threshold value of 1 applied where required. Images were exported as TIFF files for quantitative analysis.

Image quantification was performed using MetaXpress software (Molecular Devices, version 6.5.3.427). The total organoid area was segmented automatically based on Hoechst-stained nuclei, and co-localization of cell-type markers with Hoechst signal was used to quantify marker positive cells. Signal intensity was considered specific when exceeding threefold background intensity. Organoid area measurements were computed using MetaXpress internal calibration settings. Each analysis included a minimum of two technical replicates (organoids per time point) and three independent biological replicates. Each biological replicate was derived from an individual differentiation.

### Cushioned differential ultracentrifugation (CD-UC) of organoid culture conditioned media

To evaluate the Aβ profile of the secreted proteome of organoid cultures, conditioned medium (CM) was collected from the cultures at different timepoints throughout the differentiation and maturation process. Approximately 5 mL of media was removed from each of up to 6 wells of like cultures (i.e. those of the same iPSC differentiation) and these media samples were pooled and stored at -20°C. At early premature timepoints (14-18 days and 35 days) media was conditioned for 2-3 days, while at later timepoints (2.5, 4, 5, and 6 months) media was conditioned for 7 days before collection and storage.

Prior to experiments, UC bottles were sanitized with 10% hydrogen peroxide solution for at least 15 minutes. To minimize the loss of proteins to surface binding, all UC bottles, micropipette tips, and sample tubes for UC fraction storage were first blocked with a 1% BSA blocking solution; bottles, autoclaved tips, and autoclaved tubes were first filled with sterile filtered 1% BSA in 1X PBS (pH 7.4) and incubated for at least 1 hour at room temperature before being rinsed at least three times with autoclaved MilliQ water.

Frozen samples of conditioned media were thawed in room temperature water before being processed with the following cushioned differential ultracentrifugation protocol inspired by previous work by Ezparza and colleagues for postmortem tissue analyses^10^. Media was first clarified by centrifugation at 10,000 × g for 30 minutes at 4°C to remove insoluble cellular debris before the bulk portion of the clarified conditioned media (∼25 mL) was transferred into blocked ultracentrifugation (UC) bottles (Beckman Coulter cat. no. 355618). Clarified CONDITIONED MEDIA was then subjected to ultracentrifugation at 100,000 × g for 1 hour at 4°C to separate the remaining insoluble fraction, including fibrillar and protofibrillar Aβ aggregates, from smaller soluble species including peptides and oligomers. Supernatants were removed from the resulting pellets and transferred into fresh BSA-blocked UC bottles. After the transfer, ∼1 mL was removed and stored in BSA-blocked Eppendorf tubes for future analyses. Into the bottom of each UC bottle of clarified media, 1 mL of sterile filtered 70% sucrose in 1X PBS (pH 7.4) was added using an extended length blunt-tipped steel syringe needle. Cushion addition was performed with caution to avoid scaping the bottles and intermixing the solutions, as was the translocation of cushioned bottles.

Blocked UC bottles containing the CONDITIONED MEDIA and 70% sucrose cushion were then subjected to ultracentrifugation at 257,600 × g for 2 hours at 4°C. The UC bottles were then transferred to a cold room (4°C) with care to minimize agitation during transport. After clamping the bottles in place on a ring stand, 1 mL layers of the contents were carefully removed from the surface of the solution by BSA-blocked micropipette and transferred into BSA-blocked sample tubes. Generally, the upper 10 milliliters and the following 6-9 mL were pooled as two bulk upper fractions while the remaining solution including the bottommost 70% sucrose cushion was removed and stored in ∼1 mL fractions. These “cushion interface” and sucrose cushion fractions were labeled according to their proximity to the final sucrose fraction (Sucrose ”S”, S+1, S+2, etc).

### Quantification of Aβ_42_, Aβ_40_, and AD-associated soluble Aβ_42_ oligomers by ELISA

In the interest of determining the relative distribution of Aβ_42_ species of different sizes in conditioned media, Aβ_42_ and, in later experiments, AD-associated soluble Aβ_42_ oligomers were quantified in UC fractions by ELISA targeting Aβ_42_ peptide and a dodecameric variant of Aβ-derived diffusible ligand (ADDL). Additionally, portions of conditioned media were collected after the initial clarification and after the first round of ultracentrifugation to determine the concentrations and relative abundances of Aβ_42_ and Aβ_40_ by commercial ELISA.

During initial experiments aimed at determining and comparing the relative distribution of Aβ_42_ peptide across UC fractions, a sandwich ELISA against the peptide was prepared in-house as follows; 96-well ELISA plates (NUNC MaxiSorb, ThermoScientific cat. no. 442404) were prepared with 50 µL of coating antibody solution per well (coating antibody ID: Abcam, AB281143 / EPR22445) at 2 µg/mL in coating buffer (50 mM sodium bicarbonate/carbonate, pH 9.6). Coated plates were incubated sealed overnight at 4°C. Coated wells were emptied by inverting and tapping the plate onto an absorbent surface and then washed 4X with PBS (pH 7.4) before being emptied again by inversion and tapping. This washing process was repeated between each subsequent step described below. Next, 200 µL of sterile filtered 3% BSA blocking solution in PBS (pH 7.4) was added to each coated well and incubated for 2 hours at room temperature. Stock solutions of 0.1 mg/mL synthetic Aβ_1-42_ peptide (Abcam, cat. no. AB120301) were prepared in DMSO as recommended by the manufacturer, and 10 µL aliquots were stored at -80°C. Aβ_42_ standards were prepared by first diluting the DMSO stock solution to 20 ng/mL in filtered 3% BSA block solution and then performing 2X serial dilutions in 3% BSA yielding 8-11 point standard curves. After washing the blocked plates, standards and duplicate samples of 75 µL were loaded into ELISA wells using BSA-blocked micropipette tips. ELISA plates were then sealed and incubated at 37°C with shaking (∼200 rpm) for 90 minutes. Detection antibody (det antibody ID: AB280993 / EPR9296) was biotinylated according to manufacturer’s specifications (Abcam Lightning Link™ ab201795 (STN-273705, STN-273704)) and stored at 4°C. Before use, detection antibody was diluted to 0.5 µg/mL in 3% BSA blocking solution. After the standards and samples were removed and plates were washed, 75 µL of biotinylated detection antibody solution were loaded into each well before plates were sealed and incubated for 1 hour at room temperature. Streptavidin-HRP solution (Pierce cat. no. 21132) was then diluted in 3% BSA according to manufacturer’s specifications and 60-75 µL were loaded into each well. Plates were then sealed and incubated for 30 minutes at room temperature. After the final wash, 100 µL TMB solution (Thermoscientific cat. no. 34028) was loaded into each well by multi-channel pipet and incubated at room temperature protected from light until sufficient color development was achieved (5-6 minutes), at which point 100 µL of 2 M sulfuric acid was added by multichannel pipet to stop the reaction. Absorbances at 450 nm (A_450_) were then measured with a VarioSkan Lux™ microplate reader (Thermo Scientific) and Aβ_42_ concentrations were quantified with 4PL fitting of the standard curve using Skanit software (v7.0.2). After duplicate sample concentrations were averaged, the relative distribution of Aβ_42_ across cushioned UC fractions was determined by multiplying Aβ_42_ concentrations by fraction volumes and dividing each quantity by the sum total quantity of Aβ_42_ across fractions.

To quantify and determine the isolation of AD-associated soluble oligomers of Aβ_42_ in cushioned UC, representative fractions proximal to and distant from the sucrose cushion were measured by commercial ELISA against dodecameric ADDL in accordance with the manufacturer’s protocol (ThermoScientific Human Amyloid beta (Aggregated) ELISA Kit, cat. no. KHB3491). With a few exceptions, ADDL ELISAs were performed for cushioned UC pro-cessed media sourced from cultures of fully matured organoids, namely at 4–6-month timepoints. For these experiments, the uppermost and select cushion interface fractions of UC–processed media (usually S+4, S+2, and S+1) were measured to test for the expected depletion of ADDLs in the former and, if present, their enrichment in the latter. Quantification of ELISA by A_450_ was performed as described earlier.

A complementary analysis was performed to determine the quantity of Aβ_42_ and Aβ_40_ and the Aβ_42_/Aβ_40_ ratio in both conditioned media prior to CD-UC and lysates. Unlike in the previously described Aβ_42_ ELISAs focused primarily on the relative abundance of the antigen across simultaneously measured fractions, these experiments required a more robust quantitative comparison of two targets. In the interest of ensuring more robust quantitative comparison and confidence in plate-to-plate reproducibility, commercial ELISA kits were used instead of manually prepared plates. To determine and compare the concentrations of Aβ_42_ and Aβ_40_ and calculate the resulting Aβ_42_/Aβ_40_ ratios in the secretomes and organoids of all lines, samples of clarified CONDITIONED MEDIA and organoid lysates prepared by freeze/thaw cycles in RIPA buffer were analyzed by commercial ELISA kits targeting the two peptides (ThermoScientific Human Amyloid beta 42 ELISA Kit, Ultrasensitive, cat. no. KHB3544; Human Amyloid beta 42 ELISA Kit, cat. no. KHB3441; Human Amyloid beta 40 ELISA Kit, cat. no. KHB3481) according to manufacturer’s protocol. Samples were diluted to a practical concentration using the provided dilution buffer in BSA-blocked microfuge tubes. Generally, CONDITIONED MEDIA was diluted 1:2 and/or 1:4 for Aβ42 ELISA and 1:20 and/or 1:40 for Aβ40 ELISA. In cases where two levels of dilution resulted in concentrations within the range of the standard curves, concentrations were averaged before ratio calculations. Quantification of ELISAs by A_450_ was performed using the Varioskan Lux plate reader and Skanit software as described earlier. The resulting concentrations of Aβ_42_ and Aβ_40_ in pg/mL were converted to molar concentration units before calculating the molar Aβ_42:_Aβ_40_ ratio. Peptide concentrations in organoid lysates were normalized to the total protein concentration of the lysates as determined by BCA analysis according to manufacturer’s protocol (Pierce Rapid Gold BCA Kit, cat. no. A5560).

### Proteostat seed amplification assay

To assess the propensity of secreted Aβ_42_ species in UC fractions, a fluorescence-based seed amplification assay was performed. For this analysis, Proteostat (Enzo Life Sciences, cat. no. ENZ 51023) was selected as an appropriate detection reagent due to its specificity for the β-sheet tertiary structure of amyloid-type protein aggregates. For each experiment, samples were prepared in the wells of a black walled 384-well plate (µClear, Greiner) with a final volume of 40 µL containing the quantity of detection reagent recommended by the manufacturer. Seeded and non-seeded sample wells were first loaded with 20 µL of a 2X master mix solution consisting of 2X detection reagent (4% Proteostat) and 2X (10 µM) Aβ_42_ synthetic peptide from 0.1 mg/mL aliquots in DMSO (Abcam, cat. no. AB120301). Separately, positive and negative control wells were included of the same final volume containing 1X (2%) Proteostat detection reagent and 5 µM aggregated and unaggregated lysozyme controls from the assay kit. Into the non-seeded control wells, 20 µL of PBS was added to the master mix for a final volume of 40 µL and final concentration of 5 µM Aβ_42_ synthetic peptide (22.5 µg/mL). Seeded wells also contained a final concentration of 5 µM Aβ_42_ synthetic peptide and were seeded by the addition of 2-20 µL of seeding sample diluted to 20 µL with 1X PBS.

For assay mixtures seeded by UC fractions, the seeding sample volumes were adjusted based on Aβ_42_ concentrations such that approximately 2 pg of Aβ_42_ peptide was added to each seeded well. This quantity was selected to minimize the influence of the seed sample itself on initial fluorescent response. In theory, ensuring a minimal influence of seeding samples on fluorescence should allow for a consistent lag phase baseline in seeded samples whether they contain aggregate templating species, while samples with aggregate seeds would diverge during the log phase as synthetic Aβ_42_ increasingly incorporates into aggregates during templated growth. Beyond ensuring that fluorescence was primarily a result of templated aggregate growth, the 2 pg of seeding Aβ_42_ was also selected based on the Aβ_42_ concentrations in CD-UC fractions of interest. By normalizing the seed quantity to 2 pg, fractions of interest with higher concentrations could simply be diluted as needed while samples with concentrations as low as 0.1 ng/mL could be comparably analyzed by using the maximum allowed 20 µL seed volume. Seeding sample volume was limited to 50% of the final assay volume (20 µL) to allow for the standardized addition of 20 µL of 2X master mix solution to each well. All experimental samples in seed amplification experiments were prepared in duplicate. Immediately after sample preparation, plates were sealed by ELISA sticker and inserted into a preheated VarioSkan Lux™ microplate reader (Thermo Scientific) and subjected to 12-24 hours of incubation at 37°C. In 10-minute intervals during incubation, plates were shaken for 5 seconds (600 rpm, high force setting) before each reading. Amyloidal aggregate quantification was performed every 10 minutes (after/before shaking) by excitation at 550 nm immediately followed by fluorescence measurements at 600 nm.

## 3. Results

### Alzheimer’s Disease Mutations do not Alter Organoid Development

Cortical organoids recapitulated early human cortical patterning, consistent with published descriptions of inside out laminar organization in cerebral organoids^8^. The developing cortex establishes sequential neuronal layers, TBR1 and CTIP2 for deep-layer neurons, and SATB2 for upper layer neurons, while MAP2 reflects neuronal maturation. Both wild-type (WT; 7889SA New York Stem Cell Foundation) and double knock-in (DKI; BN0002, PSEN1-M146V/M146V and APP-Swedish/Swedish, New York Stem Cell Foundation) iPSC-derived organoids exhibited comparable developmental trajectories (Figure 1, Figure S1). TBR1, CTIP2, and SATB2 expression appeared sequentially and persisted over time, indicating normal temporal specification of cortical neurons in both genotypes (Figure 1). During early stages, neurons frequently co-expressed multiple cortical markers, consistent with prior observations of overlapping laminar identity during early corticogenesis^11^. The relative proportion of layer specific marker positive cells as percentage of Hoechst positive nuclei did not differ significantly between WT and DKI across all timepoints (p > 0.05) (Figure 1). These findings indicate that the Alzheimer’s associated mutations did not affect neurogenesis or layer specification during cortical development *in vitro*.

**Figure 1:**
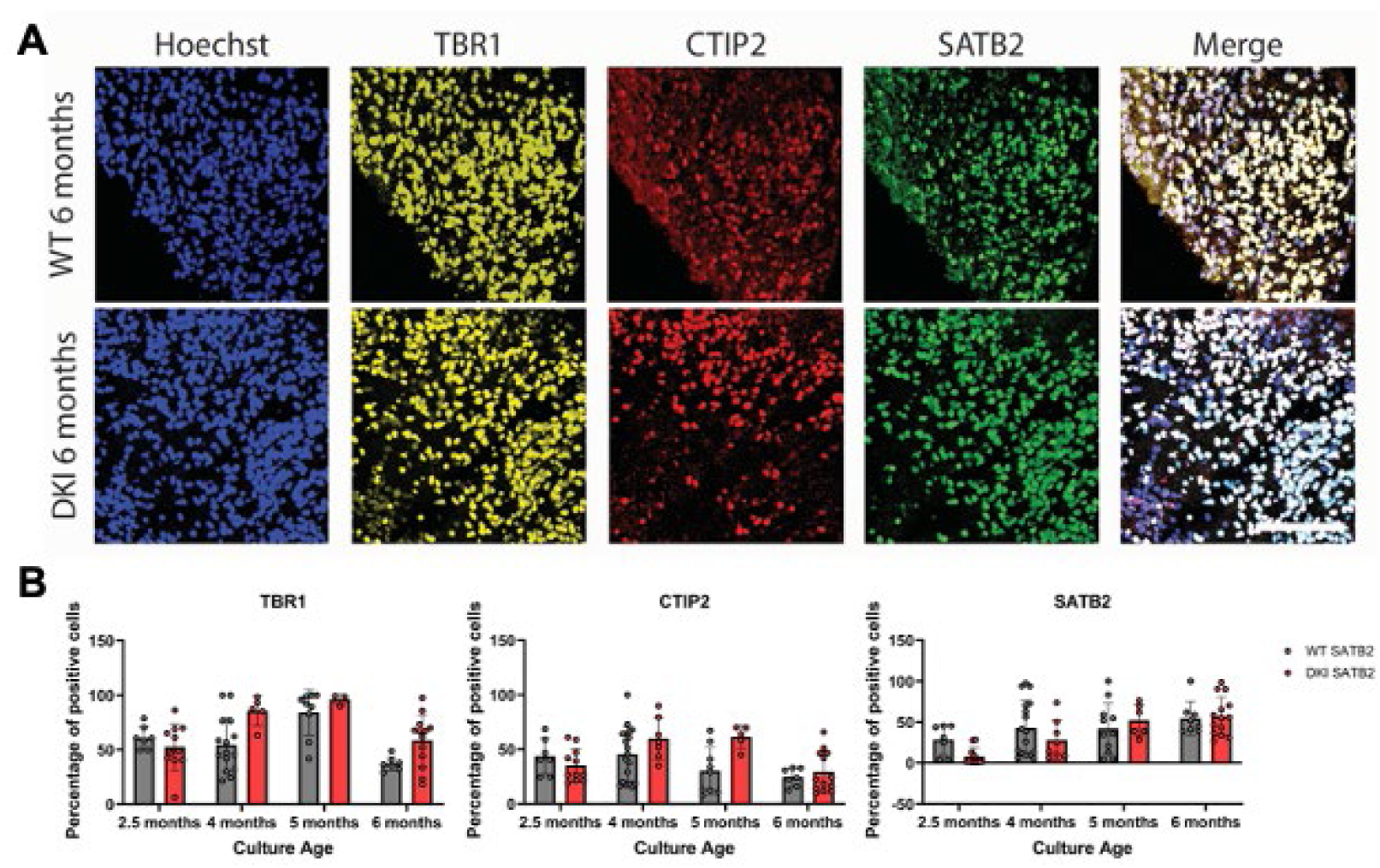
Representative image of cortical markers in cerebrocortical organoid sections. A) Images are a max projection of 15, 1-µm-thick z-stacks. Scale bar 100 µm. B) Quantification of images in **A** showing relative percentages of cortical markers; TBR1 (Layer IV), CTIP2 (Layer V), SATB2 (Layer VI) at different time points. Each point is an organoid imaged from at least 3 independent differentiations. Bars represent the mean ± SD.

### Amyloid-β Expression in Cortical Organoids

To examine amyloid-β (Aβ) pathology, organoid sections were immunostained using the 6E10 antibody (recognizing residues 1–16 of Aβ) together with either MAP2 or cortical transcription factors. MAP2 staining confirmed intact neuronal morphology and normal differentiation across all samples. Over the course of organoid development, 6E10 produced heterogeneous expression patterns in both genotypes. In some regions, 6E10 colocalized with MAP2-positive neuronal structures, following the dendritic architecture, in other regions, the signal appeared as a diffuse halo within the tissue, and occasionally it accumulated as discrete puncta or dense “blobs.” (Figure 2)

**Figure 2:**
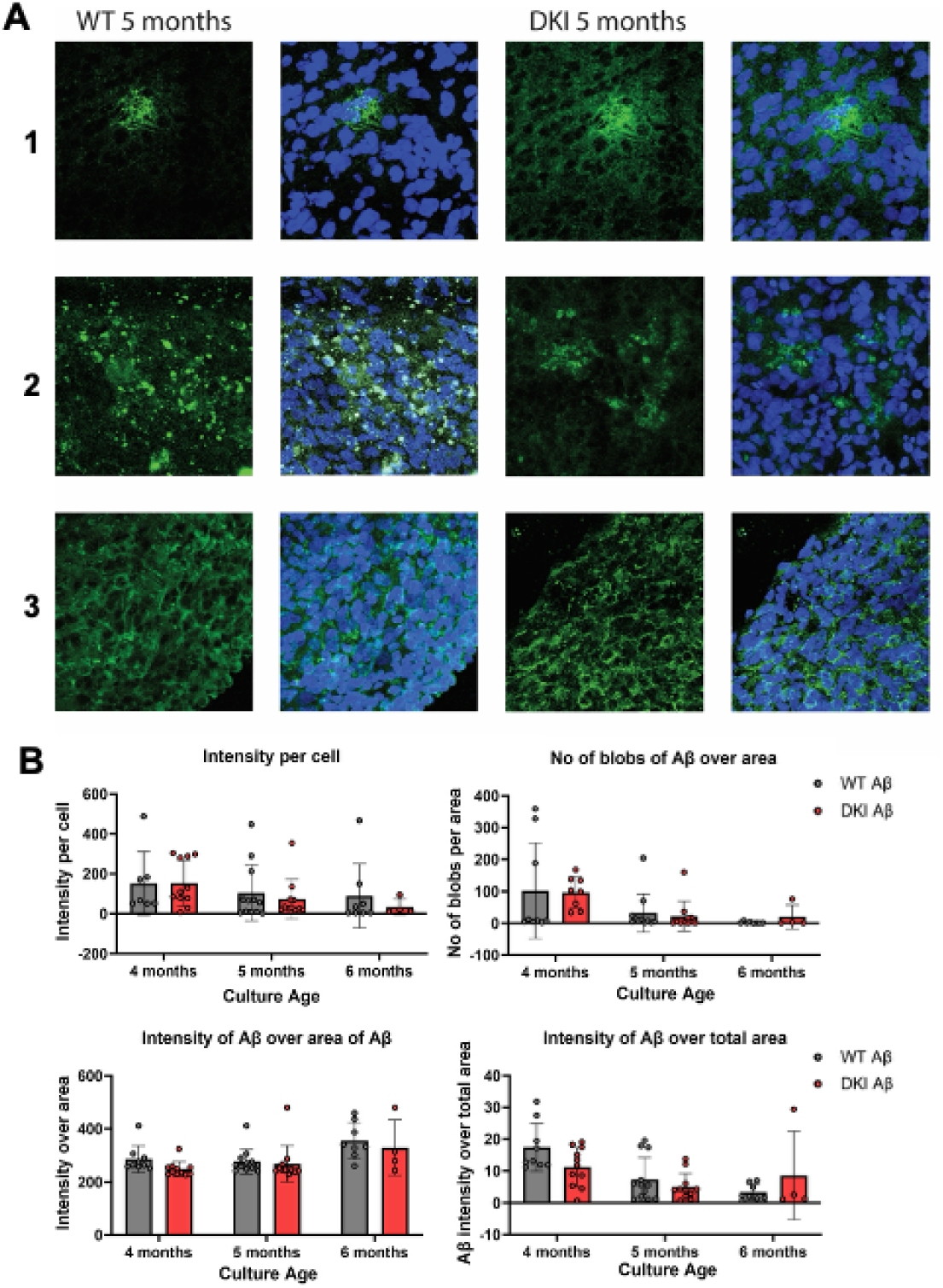
Representative image of various type of Aβ plaques observed in our cerebrocortical organoids. A) Aβ deposits are stained with 6E10 Aβ Antibody (green) and Nuclei with Hoechst (blue). Scale bar: 20 µm. 1: halo, 2: speckles, 3: Neuronal distribution. B) Quantification of Aβ distribution over 4, 5 and 6 months measured through multiple parameters showing non-significant (p>0.05) differences between WT and DKI. Each point is an organoid imaged from at least 3 independent differentiations. Bars represent the mean ± SD.

Although extracellular plaques are a classical hallmark of Alzheimer’s disease, their structural definition is not universal. Human post-mortem samples typically show compact fibrillar cores surrounded by dystrophic neurites, whereas in organoid systems, particularly at early stages, more often diffuse, small aggregates are seen rather than matured plaques^12,13^. The variability in 6E10 immunolabeling posed challenges for objective classification of Aβ morphology in our organoid system. As a result, we implemented a practical quantitative strategy based on measuring signal intensity and spatial distribution, normalized either to organoid area or to the number of Hoechst-positive nuclei.

Using this approach, we unexpectedly found that overall Aβ burden did not significantly differ between WT and DKI organoids at the examined time points (Figure 2). While the DKI line expresses pathogenic APP and PSEN1 variants associated with elevated Aβ production *in vivo*, the organoid context may limit plaque maturation or late-stage aggregation, potentially explaining the levels observed here.

It has been reported that WT organoids exposed to serum can spontaneously make Aβ plaques^14^ and our organoid protocol, chosen for its reported repeatability, uses serum during maturation. To investigate if serum is causing our WT cortical organoids to show similar Aβ burden as a highly pathogenic genotype (i.e. DKI), we differentiated these lines using a serum free protocol ^9^. Despite the absence of serum, both the WT and DKI lines again showed similar distribution of Aβ. (Figure S2)

To further validate our unexpected finding that WT and DKI have the same Aβ burden, we confirmed the specificity of the 6E10 antibody to misfolded Aβ. To address this, we used the amyloid structure dependent staining assays ProteoStat and AmyloGlo. Our findings with these stains supported the specific expression patterns seen with the 6E10 Aβ antibody and again showed that WT and DKI have the same levels of amyloid (Figure S3).

To extend our findings to a real patient genotype, we generated cortical organoids from the patient derived iPSC UCSD line (WiCell, Cell Line: UCSD241i-APP2-3). This cell line is derived from a 60-year-old female with a duplication of the amyloid precursor protein gene. Like the other two lines, the Velasco protocol yielded a large amount of variability in the neuronal differentiation success and quantity of neuron types across time. The staining pattern of Aβ in the UCSD organoids was similar to the WT and DKI line. We observed Aβ halos, blobs, and Aβ-neuronal overlap (MAP2) across different organoids within and across differentiation batches. (Figure 3)

**Figure 3:**
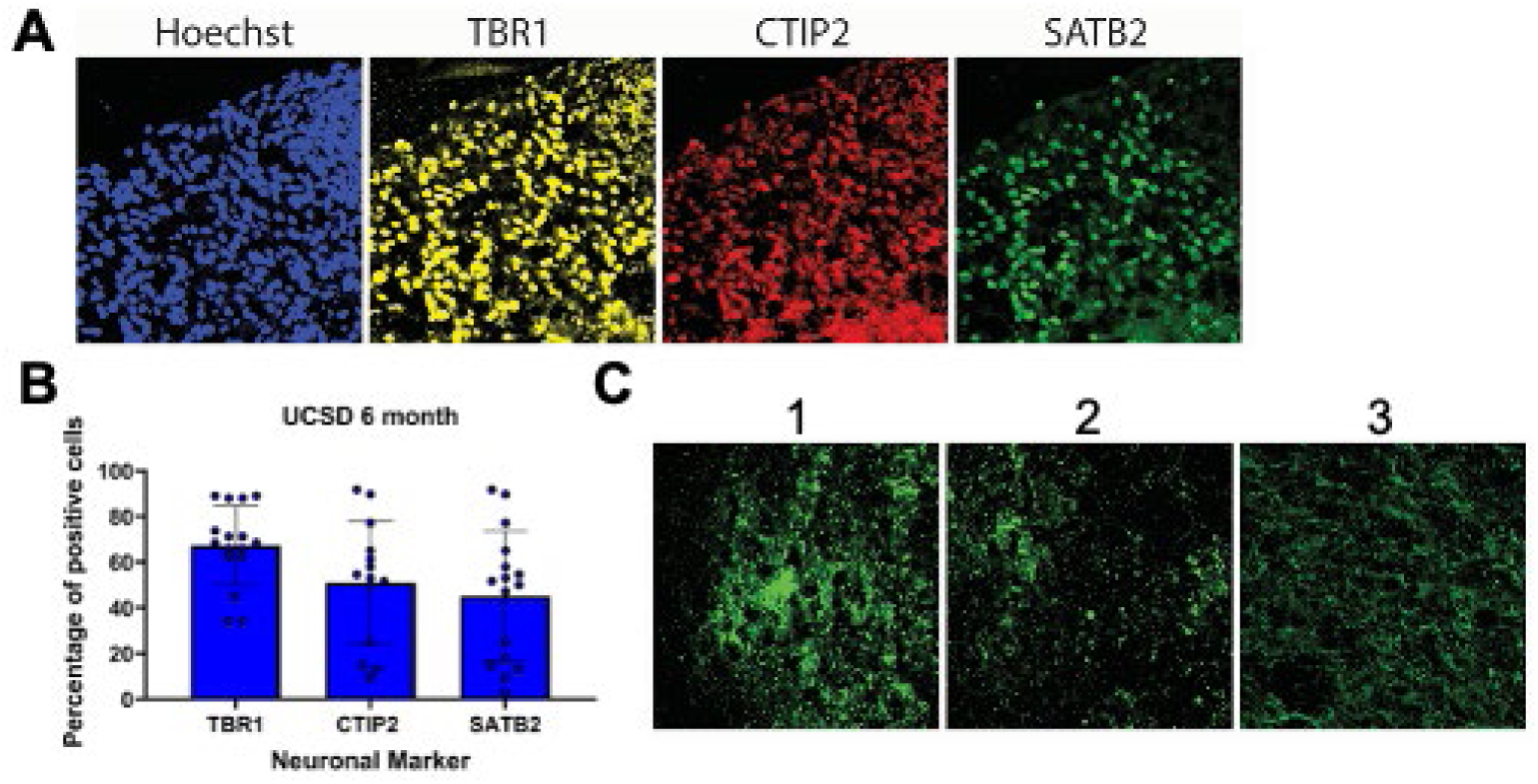
Alzheimer’s patient iPSC derived cerebrocortical organoid development and amyloid beta plaques. A) 6 month time point. Images are a max projection of 15, 1-um-thick z-stacks. Scale bar 100 µm. Quantification of images in showing relative percentages of cortical markers; TBR1 (Layer IV), CTIP2 (Layer V), SATB2 (Layer VI) at different time points. Each point is an organoid imaged from at least 3 independent differentiations. Bars represent the mean ± SD. B) Aβ deposits are stained with 6E10 Aβ antibody (green) and nuclei with Hoechst (blue). Scale bar: 20 µm. 1: halo, 2: speckles, 3: Neuronal distribution.

### Concentrations of Aβ_42_ and Aβ_40_ in Conditioned Media and Organoid Lysates

One frequent feature of familial AD (fAD) genetics is the enhanced production of Aβ_42_. While this phenotype is not universal to all fAD mutations, it is a relevant feature in the present study. Specifically, the mutations in the DKI line collaboratively contribute to increased Aβ_42_ production: APP Swe encourages amyloidogenic APP processing, increasing total β-CTF and Aβ peptide production, and PSEN1 M146V shifts γ-secretase site preference, increasing the favorability β-CTF cleavage at residue 42 and relative production of Aβ_42_. Furthermore, all APP and PSEN1 in DKI line cells are mutant forms due to the homozygous nature of the mutations. In the UCSD line, the duplication of the APP gene increases total APP production, indirectly leading to increased total Aβ_42_ through increased total Aβ.

Separate from genetic differences, several uncontrolled variables (e.g. organoid count per volume) also affect the raw Aβ_42_ concentrations in organoid culture conditioned media. While this is important to keep in mind, the starting cell densities of cultures in each line were equal, so raw Aβ concentrations in conditioned media were still compared. The relative abundance of Aβ_42_, however, is generally accepted as a more meaningful measure than its raw concentration regardless of such uncontrolled variables and it is common practice to internally normalize Aβ_42_ quantities to the more abundant, less disease-correlated peptide Aβ_40_ by calculating the Aβ_42_/Aβ_40_ ratio^1,22,23^. This superior AD biomarker is used as a diagnostic tool for AD phenotypes *in vivo* and *in vitro* with distinct interpretations in each context; *in vivo*, a reduced Aβ_42_/Aβ_40_ ratio in CSF or plasma reflects an enhanced AD phenotype as it indicates an increased sequestration of Aβ_42_ into aggregates, while *in vitro* an increased Aβ_42_/Aβ_40_ ratio reflects an enhanced AD phenotype as it indicates an increased relative production of Aβ_42_ and is further correlated with enhanced tau pathology^1,22,24^. To utilize this biomarker in the present study and determine to what degree our model aligns with previous *in vitro* results, commercial Aβ_42_ and Aβ_40_ ELISAs were used to determine the concentrations of both peptides and calculate the resulting Aβ_42_/Aβ_40_ ratios in conditioned media of mature organoid cultures. (Figure 4)

**Figure 4:**
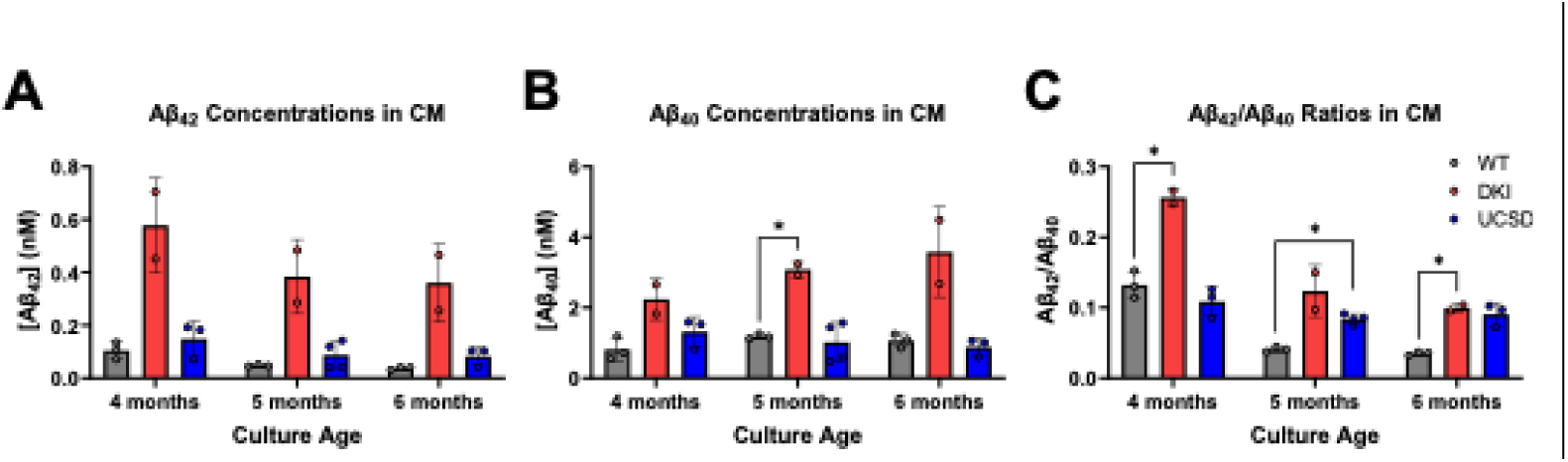
Quantification of Aβ peptide in the conditioned media of cerebrocortical organoids over time. Each point is quantification from an independent differentiation. Bars represent the mean ± SD. Each point represents an independent growth. Significant differences were determined by unpaired two-tailed t-test with Welch’s correction (*p<0.05)

In clarified WT conditioned media, average Aβ_42_ concentrations at 4-, 5-, and 6-month timepoints were 0.105±0.026, 0.048±0.003, and 0.037±0.004 nM, respectively, while average Aβ_40_ concentrations were 0.814±0.268, 1.180±0.056, and 1.063±0.161 nM, respectively, with the average Aβ_42_/Aβ_40_ ratios determined in independent growths being 0.132±0.016, 0.041±0.003, and 0.035±0.002, respectively. (Figure 4) In clarified DKI conditioned media, average Aβ_42_ concentrations at these timepoints were 0.578±0.127, 0.384±0.099, and 0.360±0.105 nM, respectively, while average Aβ_40_ concentrations were 2.239±0.415, 3.080±0.151, and 3.582±0.907 nM, respectively, with the average Aβ_42_/Aβ_40_ ratios determined in independent growths being 0.256±0.009, 0.124±0.026, and 0.100±0.004, respectively. (Figure 4) Finally, in clarified UCSD conditioned media, average Aβ_42_ concentrations at these timepoints were 0.148±0.055, 0.087±0.045, and 0.083±0.028 nM, respectively, while average Aβ_40_ concentrations were 1.314±0.346, 1.020±0.486, and 0.887±0.207 nM, respectively, with the average Aβ_42_/Aβ_40_ ratios determined in independent growths being 0.109±0.017, 0.084±0.006, and 0.090±0.013, respectively. (Figure 4)

We next wanted to compare our characterized secreted Aβ profiles to the Aβ found within the organoids. To this end, WT and DKI organoids were harvested at 6-month timepoints. Organoids were lysed in RIPA buffer, and lysates were similarly analyzed by ELISA to determine the Aβ_42_/Aβ_40_ ratios present within the organoids. The resulting ratios for WT and DKI lysates at 6 months were 0.066 and 0.087 respectively. (Figure S4)

Timepoint-matched experiments analyzing organoid lysates and conditioned media were used to compare the internal and external Aβ_42_/Aβ_40_ ratios of organoids. Specifically, two independent WT cultures and two independent DKI cultures were analyzed for Aβ_42_/Aβ_40_ ratios in both conditioned media and organoid lysates at the 6-month timepoint. In these matched experiments, WT conditioned media showed Aβ_42_/Aβ_40_ ratios of 0.036 and 0.032, or an average of 0.034, while the organoid lysates from the same cultures showed Aβ_42_/Aβ_40_ ratios of 0.060 and 0.072, or an average of 0.066. (Figure S4) In contrast, DKI conditioned media showed Aβ_42_/Aβ_40_ ratios of 0.096 and 0.104, or an average of 0.100, while the organoid lysates from the same cultures showed a Aβ_42_/Aβ_40_ ratio of 0.081 and 0.093, or an average of 0.087. (Figure S4) In this comparison, it was apparent that there was a genotype dependent difference between the secreted and organoid-bound populations of Aβ species; in WT organoids, the internal Aβ_42_/Aβ_40_ ratio was on average 93% greater than the ratio in the conditioned media, while in DKI organoids, the internal Aβ_42_/Aβ_40_ ratio was 12.97% lower than the ratio in the conditioned media. Notably, the Aβ_42_/Aβ_40_ ratios in DKI samples exceeded those of WT samples in both cases, indicating that the DKI lines produce more Aβ_42_ within the organoid and secreted into the media than the WT. In comparing the Aβ_42_/Aβ_40_ ratios further, the Aβ_42_/Aβ_40_ ratios in conditioned media of WT vs DKI organoids (WT 0.034 vs DKI 0.100) diverged to a significantly greater degree (about 2.86 times more) than the ratios in organoid lysates (WT 0.066 vs DKI 0.083, approximately 1.26 times more). This difference serves as further evidence that the secretome of AD organoid cultures is an important target for the study of AD phenotypes.

### Aβ_42_ Distributions in cushioned UC were similar in WT and DKI

Alongside the insoluble accumulation of Aβ in extracellular plaques, the extracellular secretion of soluble Aβ peptides and their self-assembly into oligomers are critical pathological hallmarks of AD^4,5,15–20^. Several variants of Aβ peptide exist, the most abundant of which are Aβ_1-X_ peptides which differ in length based on their C-terminal cleavage site and their propensity to toxically self-assemble. Among Aβ_1-X_ variants, Aβ_1-42_ or Aβ_42_ is of particular interest as it is well established to be the primary neurotoxic form with the greatest aggregability, readily seeding and incorporating into fibrillar aggregates and adopting a range of pathogenic oligomeric forms. To longitudinally investigate the secreted Aβ_42_ profile of our organoids, conditioned media samples containing the organoid secretome were collected over time for analysis. First, to determine and compare the size distribution of Aβ_42_ species in the secretomes of our WT and DKI organoids, the conditioned media was subjected to cushioned differential ultracentrifugation (CD-UC). In these experiments, soluble proteins in the conditioned media are first separated from insoluble species (e.g. fibrillar and protofibrillar aggregates) and subsequently separated from each other by density using a density cushion; larger proteins and complexes of higher density (e.g. soluble Aβ_42_ oligomers) are concentrated against a 70% sucrose cushion at the bottom of the tube while smaller, less dense species (e.g. Aβ_42_ monomers) remain in the upper fraction. Cushioned UC fractions were first analyzed by ELISA against Aβ_42_ as an initial screen for population distribution by density, and, in later experiments, by ELISA against a dodecameric Aβ-derived diffusible ligand (ADDL) to directly quantify AD-associated oligomeric species.

The conditioned media collected from different lines and independent cultures were timepoint-matched and conditioned for equal periods at each timepoint. However, while the starting cell density of differentiations were all the same, matured samples were not controlled for cell or organoid counts per volume. Due to these uncontrolled variables, Aβ_42_ distributions across fractions were calculated and compared in terms of the fraction of the total quantity of Aβ_42_ present in each processed sample. For distribution analysis, fractions were grouped by depth to delineate the regions expected to be enriched with and depleted of oligomeric Aβ_42_; the presumably enriched “cushion set” consists of the 1 mL sucrose cushion and 4-5 mL of solution in the media/cushion interface atop it, together accounting for 20-24% sample volume, while the presumably depleted “upper set” includes the remaining fractions above the former, accounting for the remaining 76-80% sample volume. Given this partitioning, Aβ_42_ enrichment in the cushion set would therefore be roughly signified by the fraction of total Aβ_42_ ± standard deviation exceeding 22%. The relative fraction of Aβ_42_ in the cushion set varied with the age of cultures similarly in both the WT and DKI lines. At the earliest studied timepoint (14-18 days), cushion sets showed apparent Aβ_42_ enrichment in both WT and DKI lines with WT cushion sets containing an average of 47.2 ± 12.8% of total Aβ_42_ and DKI cushion sets containing an average of 51.8 ± 8.9% of total Aβ_42_. (Figure 5) Interestingly, expression of Aβ peptide has been linked to normal human brain development and as such, we attribute this early timepoint Aβ_42_ enrichment to normal development and do not consider it an accurate representation of the secreted proteome of neuronal organoids^21^.

**Figure 5.**
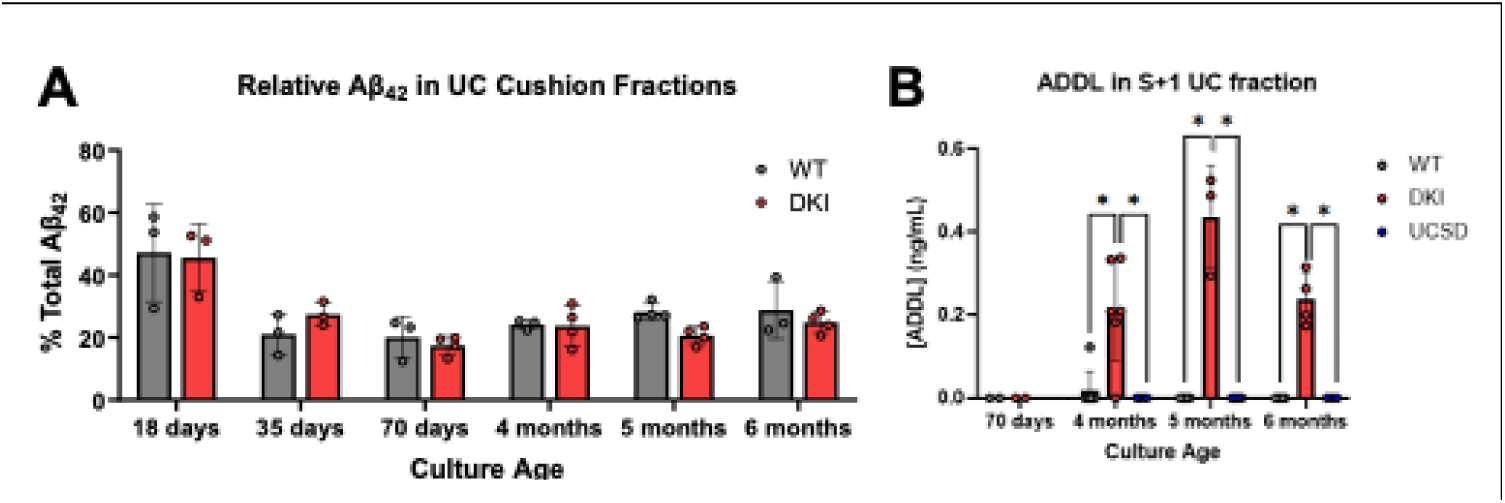
Aβ_42_ distributions in WT and DKI UC cushion fractions and ADDL concentrations in S+1 fractions. A) Percent of total Aβ_42_ present in the cushion set of UC fractions. Enrichment of Aβ_42_ is indicated by the percent of total Aβ_42_ exceeding the percent of total sample volume. Each bar represents 3-4 independent growths. B) ADDL content of the lowest (S+1) cushion interface UC fractions. ADDLs were not detected in any upper fractions *(not depicted).* Each bar represents 2-7 independent growths. Significant differences were determined by unpaired two-tailed t-test with Welch’s correction (*p<0.05)

At subsequent timepoints, the relative fraction of Aβ_42_ in cushion sets did not indicate notable enrichment in either WT or DKI lines. At day 35, day 70, 4-, 5-, and 6-month timepoints, WT cushion sets contained an average of 21.1 ± 5.3%, 20.1 ± 5.5%, 24.2 ± 1.3%, 28.2 ± 2.3%, and 28.8 ± 7.5% of total Aβ_42_, respectively. At these timepoints, DKI cushion sets contained an average of 27.4 ± 3.1%, 17.4 ± 2.6%, 23.8 ± 5.5%, 20.6 ± 2.5%, and 24.9±3.0% of total Aβ_42_, respectively. (Figure 5)

### Aβ-derived diffusible ligands (ADDL) were isolated by CD-UC consistently in the DKI derived organoids

Next, we expanded our efforts to detect and temporally map Aβ_42_ oligomer generation and secretion during organoid development. Following the determination of the relative distribution of Aβ_42_ peptide across UC fractions over time, AD-associated oligomers were directly measured in UC fractions by ELISA against a dodecameric variant of Aβ-derived diffusible ligand (ADDL). ADDLs are a heterogeneous subclass of globular Aβ_42_ oligomers that have been previously described as the most potent neurotoxic species with significant evidence implicating dodecamer bioactivity in AD pathophysiology^16–20^.

To determine the timeline for Aβ oligomer generation, conditioned media from WT and DKI cultures from the less mature day 70 timepoint and from mature organoids (i.e. 4, 5, and 6-month old cultures) was subjected to CD-UC and fractionated as described. At day 70, oligomers were not detected in any fractions of either WT or DKI conditioned media (Figure 5). However, at the mature timepoints, ADDLs were detected in cushion interface fractions of nearly all replicates of DKI conditioned media. In contrast, ADDLs were not detected in WT conditioned media fractions with the exception of a single outlier. (Figure 5) This suggests that maturation of organoids to a later stage of development, where sufficient cells are SATB2 positive, is integral to the production of secreted oligomers in either cell line. (Figure 1)

Validating our CD-UC approach to oligomer isolation, ADDLs were not detected in the uppermost fraction in any of these experiments and, whenever detected in multiple cushion interface fractions, ADDL concentrations increased with proximity to the sucrose cushion. These results therefore confirmed: that disease-relevant soluble Aβ_42_ oligomers are generated and secreted by DKI line organoids, that they are not similarly present in the conditioned media of our WT line organoids, and finally that the CD-UC protocol successfully isolates bioactive ADDLs from the bulk conditioned media of AD organoid cultures.

We considered the possibility that the ADDLs may have been present in the WT fractions in a similar proportion to the total Aβ_42_ population as observed in DKI samples, but due to the lesser total Aβ_42_ measured in WT samples, they may have been below the limit of detection by ELISA. To address this possibility, we first determined the ratio of ADDL to Aβ_42_ concentration across all DKI cushion interface S+1 fractions. To determine if Aβ_42_ concentrations in WT samples were sufficiently high for detectable ADDL assuming similar relative populations, we used both the median (0.273) and average (0.521) ADDL/Aβ_42_ ratios in the DKI fractions to calculate a range of “expected” ADDL concentrations in WT S+1 fractions. According to these calculations, 66.7-75.0% of the WT fractions contained sufficient Aβ_42_ for detectable ADDL at these median and mean ratios. These calculations confirmed that ADDLs were not present in a similar ratio in WT samples and suggest that ADDL formation in the DKI conditioned media is driven by AD genetics/phenotype rather than by Aβ_42_ concentration alone.

In addition to the WT and DKI lines, conditioned media samples collected from the patient-derived UCSD line were also processed by CD-UC and similarly analyzed by ADDL ELISA. At the 4-, 5-, and 6-month timepoints, tested cushion interface fractions did not contain a detectible quantity of ADDL oligomers.

### Seed amplification assay

Some forms of oligomeric Aβ can act as aggregate seeds that template amyloidal misfolding and recruit smaller soluble species over time into a growing fibrillar aggregate^25^. To test the aggregate seeding propensity of Aβ oligomers isolated in the cushioned UC fractions of condition media, we performed a seed amplification assay using Proteostat (Enzo Life Sciences), a dye that becomes fluorescent upon intercalation into amyloid fibrillar aggregates. In this assay, samples of UC fractions were added to a solution containing synthetic Aβ_42_ peptide and Proteostat dye and the solutions were subjected to heating (37°C) and shaking (600 rpm) over time (12-24 hours) in a plate reader with fluorescence readings throughout the duration. In these experiments, amyloidal aggregates are seeded by the UC fractions and the recruitment of synthetic Aβ_42_ into the aggregates over time leads to increased Proteostat binding and fluorescence. In the resulting sigmoidal fluorescence curves, the duration of the lag phase indicates how rapidly peptide recruitment is initiated while the maximum magnitude of the slope in the log phase indicates the peak rate of aggregate elongation. In each experiment, PBS only non-seeded controls were included for comparison and normalization.

During the analysis of the seed amplification assay fluorescence curves, we first noticed a correlation between UC fraction depth and Aβ_42_ aggregation kinetics. Seeding fractions yielded earlier nucleation and more rapid elongation with increasing proximity to the sucrose cushion. (Figure S5) This was the expected result for DKI fractions since the lower cushion interface fractions were measured to have increasingly high concentrations of ADDLs and the presence of Aβ oligomers was expected to correlate with aggregate seeding. Surprisingly, however, this trend was observed independent of genotype and, consequently, the presence of Aβ oligomers. (Figure S5) When comparing assay mixtures seeded by matched-depth WT fractions, we instead noticed a correlation between increased apparent seed amplification and the volume of UC fraction added as the seeding sample. Seeding sample volumes had been adjusted to standardize the dose of Aβ_42_ added to the peptide/dye mixture, and samples seeded by greater volumes of each UC fraction consistently yielded fluorescence curves indicating greater seeded aggregation. These results suggested that the background solution of UC-processed conditioned media fractions (i.e. the components of the organoid cortical differentiation media, or CDM) and/or secreted biomolecules other than Aβ species were contributing significantly to aggregate seeding and Proteostat fluorescence.

To test the influence of UC-processed CDM alone, fresh differentiation media was subjected to the same CD-UC protocol and fractionation, and the fractions were used in the same Proteostat seed amplification assay. In these experiments, samples seeded by the UC fractions of fresh media similarly showed increased aggregate seeding with fraction depth. Resulting fluorescence curves were indistinguishable from those of samples seeded by similar volumes of WT or DKI conditioned media fractions. (Figure S5) These results confirmed that media components in UC fractions rather than biomaterials from organoid cultures were the primary contributors to the observed aggregate seeding, and that background influences prevent the specific determination of the seeding propensity of culture-sourced Aβ species.

Following these observations, individual media components were similarly tested for aggregate seeding as well as for direct Proteostat fluorescent activation. To identify the major contributors to the influence of the media background, media components were added to Proteostat solutions at the maximum permitted seeding sample volume (i.e. 20 µL seed into 20 µL 2X Proteostat) in the presence and absence of synthetic Aβ_42_ peptide to separately determine their Proteostat binding and Aβ_42_ aggregate seeding properties. Since the previous tests showed greater seeding by lower UC fractions which contained a disproportionate concentration of dense molecular species, and the degree to which each media component was concentrated into such fractions was unclear, seeding samples were tested at their maximum/stock concentrations to measure their maximum possible influence. From these experiments, several medium components were found to encourage Aβ_42_ aggregation and bind Proteostat in isolation. In approximately descending order, the commercial components that caused the most notable aggregation of Aβ_42_ were N2 > FBS > KSR > B27 ≥ CDL with each of these components except for CDL yielding a notable Proteostat response in the absence of Aβ_42_ in the same order. (Figure S5) From these results, it was concluded that the accurate assessment of the seeding propensity of Aβ_42_ species isolated by cushioned UC could not be conducted without further extracting Aβ_42_ away from the co-isolated medium components.

## 4. Discussion

Brain organoids have been increasingly employed as a disease-modeling system due to their relative accessibility and their ability to capture the complex cellular environment of the human brain. There are many protocols available; each has itsown strengths and weaknesses, but most are shown to reliably recapitulate neuronal differentiation and development that resembles human brain development. We adopted the Velasco et al. protocol because it was reported to yield low variability between organoids, supported by RNA-seq analyses, and was shown to work with multiple cell lines^8^.

However, we did not observe such reproducibility in our cell lines. We repeatedly observed large variation in neuronal composition between organoids within a differentiation batch and across independent differentiations. Similar protocol dependent and cell-line dependent variations are also noted by other researchers suggesting that even low-variability protocols can still cause high variability depending on handling conditions^26,27^. While heterogeneous in nature, we were able to produce and maintain cortico-organoids expressing the canonical cortical markers TBR1, SATB2, and CTIP2 and these “verified” cortical organoids were used in downstream experiments. The addition of Alzheimer’s mutations did not cause any significant differences in corticogenesis and the AD lines had the same heterogeneity as the WT control. (Figure 1, 3) This aligns with human biology where the patients with these mutations do not have any symptoms until late in life^28,29^.

Unexpectedly, we consistently measured comparable levels of Aβ in the WT and DKI cortical organoids. Although the two cell lines contain the same genetic background, DKI has knock-in APP and PSEN2 mutations, both of which are commonly associated with increased Aβ. However, even after rigorous verification using independent techniques and different analytical methods, we did not observe the elevated levels of Aβ accumulation as previously seen in neuronal cultures ^30^. This discrepancy suggests that perhaps the three-dimensional context of organoids may not be sufficient to produce mature brain context for mutation-driven amyloid pathology to diverge within the tissue itself.

One possible methodological explanation for this result may be serum exposure. Others have reported that serum exposure can induce Aβ aggregation in cortical organoids^14^. As our chosen differentiation method contains serum in the maturation media, we performed additional differentiations using a serum free protocol^9^. Despite removing the serum from the differentiation, both the WT and DKI diseased line showed comparable Aβ burden within the organoids. This suggests that the lack of significant difference in our model is unlikely to be driven by serum exposure.

No distinction in Aβ accumulation between the WT and DKI could also be due to several biological factors. Cortical organoids transcriptionally resemble fetal human brain tissue and therefore may not reach the developmental or biological age required to reveal genotype-specific differences characteristic of late-onset pathology^31^ Further, AD pathology depends on interactions among neurons, vasculature, and particularly microglia, which play a critical role in plaque compaction and the transition from diffuse Aβ aggregates to mature fibrillar plaques. In our organoids, in the absence of microglia, Aβ may remain in non-specific, heterogeneous forms, appearing as puncta, diffuse halos, or neurite-associated signal, rather than forming organized plaques.

Additionally, the 6E10 antibody recognizes soluble and insoluble Aβ species as well as full-length APP, making it difficult to distinguish between pathogenic aggregates, early intracellular intermediates, and baseline APP expression. However, we verified our results by performing ProteoStat and Amyloglo assays which support that what we are measuring is amyloid in structure.

Technical factors also added to the complexity of interpretation of amyloid beta plaque burden as there is no universally accepted definition of a plaque, either in terms of size, morphology, or distribution pattern. Indeed, all types of distributions of plaques we observed in our study have been reported in the literature^32–34^. However, the absence of a defined system for characterizing plaques adds challenges to consistently analyze the burden and compare across different studies.

Finally, our findings could also align with population-level observations that Aβ plaques are found in a substantial proportion of cognitively normal individuals. Only the DKI mutant line demonstrates the capacity to generate soluble, disease-relevant oligomers (ADDLs), suggesting that the AD mutations specifically drive the formation of neurotoxic species rather than simply increasing overall amyloid accumulation.

As the extracellular soluble oligomers of Aβ are thought to be the toxic species^4,5,15^, we wanted to understand when in organoid development do Aβ peptides begin to be secreted, do they oligomerize, and do we see an AD-like Aβ_42_ to Aβ_40_ ratio in our mutant lines. To this end, the conditioned media from WT and the AD lines DKI and UCSD were analyzed throughout their development via ELISA. While lacking statistical significance, there were obvious trends in the Aβ_42_ and Aβ_40_ concentrations and ratios. The concentrations of secreted Aβ_42_ and Aβ_40_ in DKI conditioned media were consistently greater than WT control at each mature timepoint (4-, 5-, 6-month), and timepoint-matched comparisons also showed elevated Aβ_42_/Aβ_40_ ratios versus WT control. Considering that the fAD mutations in the DKI line (APP, PSEN2) increase the total production of Aβ and Aβ_42_ *in vivo*, this result aligned well with expectations. In the conditioned media of the patient-derived UCSD line organoids, the relative concentrations and ratios of Aβ_42_ and Aβ_40_ were more consistent across mature timepoints than either WT or DKI. Across mature timepoints, UCSD conditioned media Aβ_42_/Aβ_40_ ratios approximately aligned with those of DKI media at 6 months with little variation to raw concentrations.

It is worth noting that while DKI consistently had higher Aβ_42_/Aβ_40_ at each timepoint, the ratios changed from month to month, seemingly decreasing over time. This indicates that establishing a universal/definitive Aβ_42_/Aβ_40_ threshold indicative of AD pathology in the organoid secretome is nontrivial, and this emphasizes the importance of timepoint-matched comparisons. While tests for Aβ_42_/Aβ_40_ ratios in CSF and blood plasma have well established correlations to AD risk, Aβ_42_/Aβ_40_ ratios in human cell culture models vary significantly across model types (monocultures vs. cocultures, 2D vs. 3D) and differentiation protocols^1^.

Age-dependent increases in extracellular amyloid plaque deposition have been reported in previous AD organoid studies, recapitulating the progressive growth of amyloid plaques in the aging human brain^2^. Intuitively, an age-dependent decrease in the relative abundance of soluble Aβ_42_ in the conditioned media could be attributed to the increased accumulation of Aβ_42_ into extracellular plaques over time. However, Aβ staining in sectioned organoids did not indicate an age-dependent increase in Aβ plaques by any measure but instead showed a reduction with age. Alternatively, the measured decrease of secreted Aβ may be from increasing amounts of insoluble protofibrillar and/or fibrillar aggregates in solution being removed along with cellular debris in the first step of conditioned media collection.

Our initial approach to quantifying the degree of Aβ oligomerization in conditioned media was to establish the population distribution of Aβ_42_ species by density by measuring Aβ_42_ peptide across cushioned UC fractions. In these analyses, we found that relative distributions of Aβ_42_ peptide across UC fractions of conditioned media did not clearly indicate significant cushion interface enrichment past premature timepoints in either WT or DKI lines. However, Aβ_42_ oligomers have varying degrees of peptide burial and assembly stability, so the use of antibodies against Aβ_42_ peptide may fail to accurately capture the true total abundance of the peptide in oligomer-enriched fractions due to epitope obstruction and/or in situ disassembly after antibody capture. Because of this, the apparent lack of cushion interface enrichment by these measures was not necessarily conclusive, so we instead proceeded to directly measure an Aβ-derived diffusible ligand epitope by ELISA. With this approach, no upper fractions contained detectable ADDLs while cushion interface fractions of nearly all the DKI samples and nearly none of the WT samples contained ADDLs with a single exception for each. Although the patient-derived UCSD line did not yield detectable quantities of the targeted ADDL in UC isolates, this does not necessarily imply that the fractions are devoid of all AD-relevant oligomers.

ADDLs in DKI cushion interface fractions were low abundance with concentrations in the range of hundreds of pg/mL. These isolates were prepared from samples generated in a short timeline by 7 days of media conditioning in 6 well plate cultures. By maintaining a larger number of organoid cultures and collecting and processing conditioned media at every feeding, this approach can be scaled up and leveraged to produce and isolate AD-associated bioactive AβOs secreted from a genotype-specific physiologically relevant source.

Although fluorescence-based seed amplification experiments were intended to investigate the amyloidal aggregate seeding propensity of UC-isolated Aβ populations, the discovery of significant background influences on Aβ fibrilization and Proteostat fluorescence by co-isolated media components obstructed these analyses. While the results of these experiments did not produce useful data for their original intended goal, they still provided insights that permitted some important inferences. First, several media components, primarily N2 > FBS > KSR > B27 ≥ CDL were found to notably contribute to Aβ fibrilization and/or direct Proteostat response. The presence of any of these materials, especially if their concentrations in tested samples are independently varied, is therefore an important consideration in studies aiming to quantify the β-sheet tertiary structure of specific amyloidal aggregates by similar fluorescent dyes. While these components in UC fractions were observed to seed Aβ fibrilization, it is important to note that their presence did not similarly influence the formation of ADDLs. Specifically, the media components were similarly co-isolated into cushion interface fractions during cushioned UC of conditioned media from all lines, but ADDLs were only consistently detected in DKI fractions. This indicates that, while the media components may encourage amyloidal proto-fibril or fibrillar aggregate assembly, they do not similarly encourage the generation of the off-pathway globular ADDL species of greater AD relevance.

Despite an increasing shift in focus from the amyloid aggregate hypothesis towards the amyloid oligomer hypothesis, studies focused on the production/presence of soluble oligomers in tissue culture conditioned fluids are surprisingly scarce^4,5,15^. It is particularly surprising that few studies have attempted to isolate and structurally characterize the soluble oligomeric Aβ species that are actively secreted into solution by disease-relevant tissues. Although Aβ oligomers are highly heterogeneous and remain poorly characterized, the amyloid oligomer hypothesis suggests that a subset of these species is a major driving force, possibly the primary driving force, of AD pathology. Aβ oligomers begin to form in early pre-symptomatic stages of disease progression and exert neurotoxic effects through several direct and indirect mechanisms with extracellular diffusible species propagating AD pathology and seeding amyloid aggregation^4,5,15,19,35^. Among Aβ oligomers, a highly toxic globular subset of soluble species called Aβ-derived diffusible ligands (ADDLs) are of key interest and can be found in both the CSF and the water-soluble fractions of AD brain homogenates as well as cultured neuronal extracts. The Aβ oligomers in lysates and homogenates, however, broadly include a mixture of species released from membranes, extracellular vesicles, the intracellular space, and fragmented plaques, as well as those assembled following the intermixing of these forms with free peptides. Since some degree of perturbation to the soluble populations and oligomerization in tissue extracts is unavoidable, it is difficult to resolve with certainty which forms in the soluble milieu are natively assembled, active participants in pathogenic mechanisms prior to homogenization^15^. We therefore posit that in the present study and future studies employing similar methods, the disease relevance and mimicry of early-stage *in vivo* conditions is further optimized and preserved by instead focusing on the isolation, quantification, and characterization of Aβoligomers present in the soluble secreted proteome of AD brain organoids. The generation and secretion of soluble AβOs by AD organoids provides a unique, sparsely explored opportunity to directly study from the secretome with minimal perturbation to their natively folded state, and these analyses are especially physiologically relevant when utilizing patient-derived iPSC-derived cerebrocortical organoids models given their cellularly diversity and brain-like layered structure.

## 5. Conclusions

The advent of iPSC-derived cerebrocortical organoids, tissue cultures that recapitulate the complex cytoarchitecture of the human brain, has brought about a new era of physiological relevance in the *in vitro* modelling of neurodegenerative diseases. With recent developments in our understanding of the protein pathologies of Alzheimer’s disease (AD), an increased focus has been placed on the highly toxic soluble oligomers of amyloid beta (Aβ) secreted by the AD brain rather than their deposition into plaques. Despite the shift in interest from Aβ plaques to Aβ oligomers, or AβOs, these heterogeneous assemblies remain insufficiently characterized. While greater quantities of various AβOs have been detected in homogenates of AD brains and cultured neuronal tissue, the homogenization process inherently influences the oligomerization and populations of soluble Aβ. Protein aggregation is progressive, and oligomer population distributions fluctuate, so while homogenization is far from the only source of *ex vivo / ex vitro* artifacts that complicate oligomer studies, efforts to minimize such artifacts are critical. The secreted proteome, or secretome, of iPSC-derived AD cerebrocortical organoids offers a uniquely physiologically relevant source for secreted, natively assembled soluble AβOs implicated in AD pathophysiology. By isolating secreted AβOs directly from this soluble reservoir, tissue homogenization artifacts are circumvented, and the preservation of extracellular AβO population distributions is improved.

the present study, cerebrocortical organoids were successfully derived from iPSCs of isogenic AD and WT organoids as well as a patient derived fAD line. Imaging of organoid sections throughout development confirmed their neuronal differentiation and cytoarchitecture however intratissue Aβ levels did not significantly differ, with all lines showing similar Aβ plaque burden. While Aβ plaque burden is a traditional measure of AD phenotype, a substantial fraction of healthy humans has significant amyloid plaques^7^. Our results add to the growing evidence that this metric is insufficient to identify AD phenotypes, even in highly aggressive Aβ-enhancing genotypes. In contrast, mature cerebrocortical organoids derived from the isogenic lines yield conditioned media containing distinct secreted Aβ profiles. According to ELISA analyses, the secretome of mature DKI organoids consistently showed elevated levels of Aβ_42_ and Aβ_40_, and elevated Aβ_42_/Aβ_40_ ratios compared to the WT line at all timepoints in alignment with the expected AD phenotype. After subjecting conditioned media to cushioned differential ultracentrifugation, the presence and successful isolation of AD-associated Aβ-derived diffusible ligands was consistently confirmed exclusively in DKI samples. With this novel approach to isolating bioactive species from the secreted proteome of AD brain organoids, we present a promising advancement in the study of soluble oligomeric Aβ.

It is our opinion that this study justifies an increased focus on the secreted proteome in the conditioned media of brain organoids in Alzheimer’s disease research, both as an underutilized aspect of the model in defining AD phenotypes, and as a physiologically relevant scalable source for bioactive Aβ oligomers. Pairing this approach with further AβO purification (e.g. non-denaturing immunoprecipitation) and subsequent structural analyses (e.g. circular dichroism, electron microscopy, atomic force microscopy) and neurotoxicity assessments (e.g. application to healthy neurons) could potentially contribute significantly to ongoing efforts to characterize early-stage AD-causative species. There are several ways that future studies can further expand on the protocol to investigate additional variables to AβO formation. For example, the isolation of AβOs from non-oligomeric Aβ in solution could be leveraged to investigate post-translational modifications to Aβ that occur with greater frequency in oligomer enriched fractions, and patient-derived iPSCs and engineered fAD genetic knockouts could be used to delineate the upstream influence of distinct fAD genetics on Aβ oligomerization. By expanding on this work with such future studies, a novel route to designing patient-tailored treatments could be explored by narrowing the scope of AβO-targeting therapeutics to those species identified in conditioned media isolates of gene- or patient-specific organoid cultures.

## Supplementary Materials

Figure S1: Organoid development over time; Figure S2: Serum-free protocol examples of amyloid beta plaques; Figure S3: Proteostat and AmyloGlo Images; Figure S4: Quantification of amyloid beta in brain organoid lysates; Figure S5: Amyloid seed amplification assays

## Author Contributions

Conceptualization, A.O.; methodology, E.Z.; formal analysis, A.O., E.Z., M.F.; investigation, E.Z., M.F., S.B.; resources, A.O.; data curation, A.O., E.Z., M.F.; writing—original draft preparation, E.Z. M.F.; writing—review and editing, A.O., E.Z., M.F., S.B.; supervision, A.O.; project administration, A.O.; funding acquisition, A.O. All authors have read and agreed to the published version of the manuscript.

## Funding

Research reported in this publication was supported by the National Institute On Aging of the National Institutes of Health under Award Number R01AG078187. The content is solely the responsibility of the authors and does not necessarily represent the official views of the National Institutes of Health

## Acknowledgments

We would like to thank Dosh Whye (Assistant Director, Human Neuron Core, Boston Children’s Hospital) for his insightful input and help in developing our brain organoid differentiation protocols.

## Conflicts of Interest

The authors declare no conflicts of interest

## Abbreviations

The following abbreviations are used in this manuscript:

AD: Alzheimer’s Disease
Aβ: Amyloid beta
AβOs: Amyloid beta oligomers
UC: Ultra centrifugation
WT: Wild-type
DKI: Double knock in
ADDLs: Aβ-derived diffusible ligands

## Notes

### Competing Interest Statement

The authors have declared no competing interest.

