## Supplemental Information for "Alzheimer’s Disease Brain Organoids as a Source of Disease-Relevant Amyloid-Beta Oligomers"

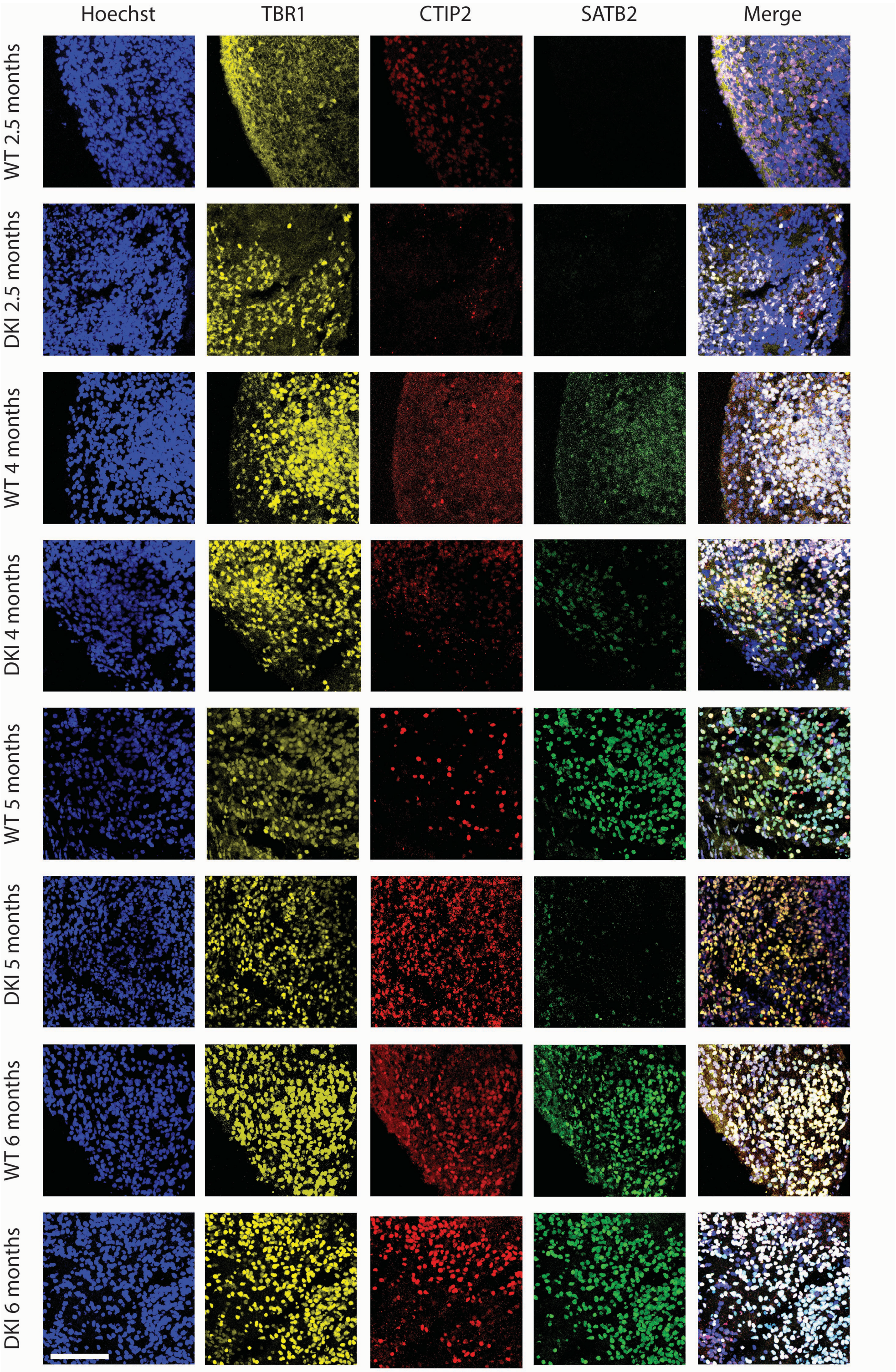


**Supplemental Figure 1: Tracking cerebrocortical organoid development over time.** Representative image of cortical markers in WT and DKI cerebrocortical organoid sections over time (2.5, 4, 5 and 6 months age) Images are max projections of 15 1-μm-thick z-stacks. Scale bar: 100 μm.

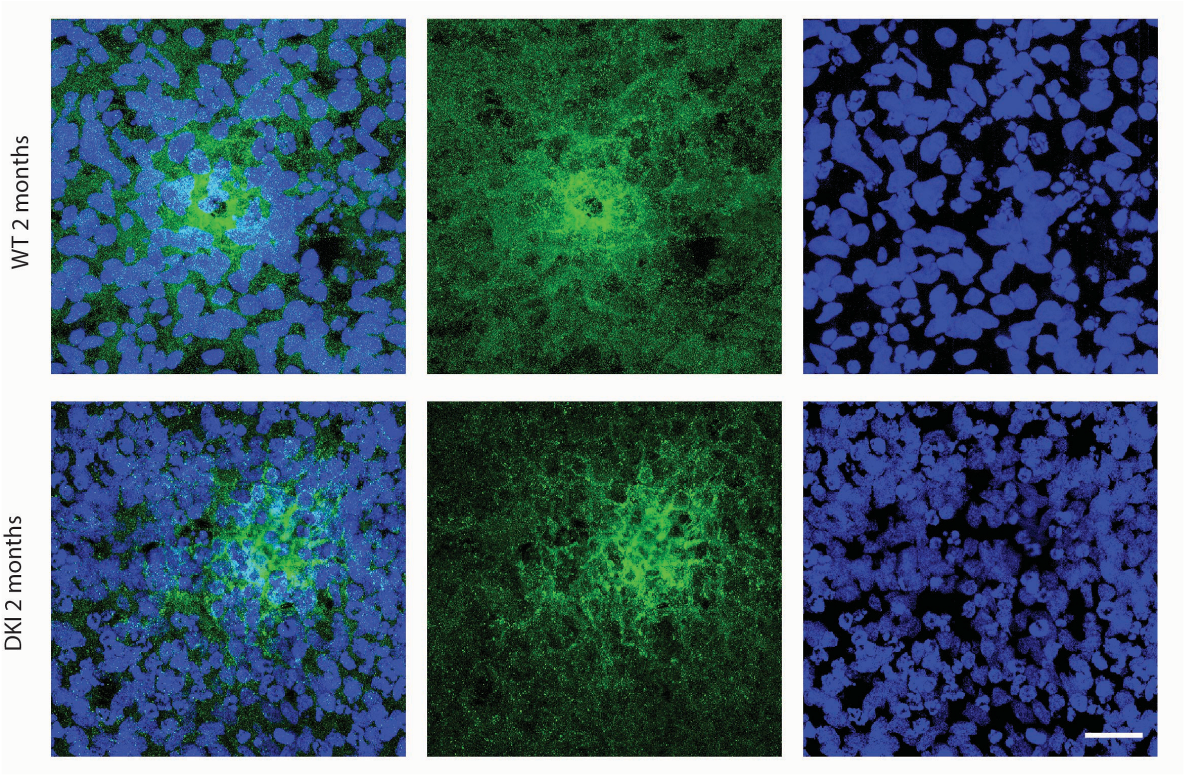


**Supplemental Figure 2: Representative images of amyloid-beta plaques seen in cerebrocortical organoids differentiated using a serum free protocol** (Whye, et al., Curr Protoc 3(1) 2023). Hoechst (nuclei, blue) anti- amyloid-beta antibody 6E10 (green). Diffuse plaques are seen at 2 months in both the WT and DKI derived cerebrocortical organoids. Scale bar 20μm.

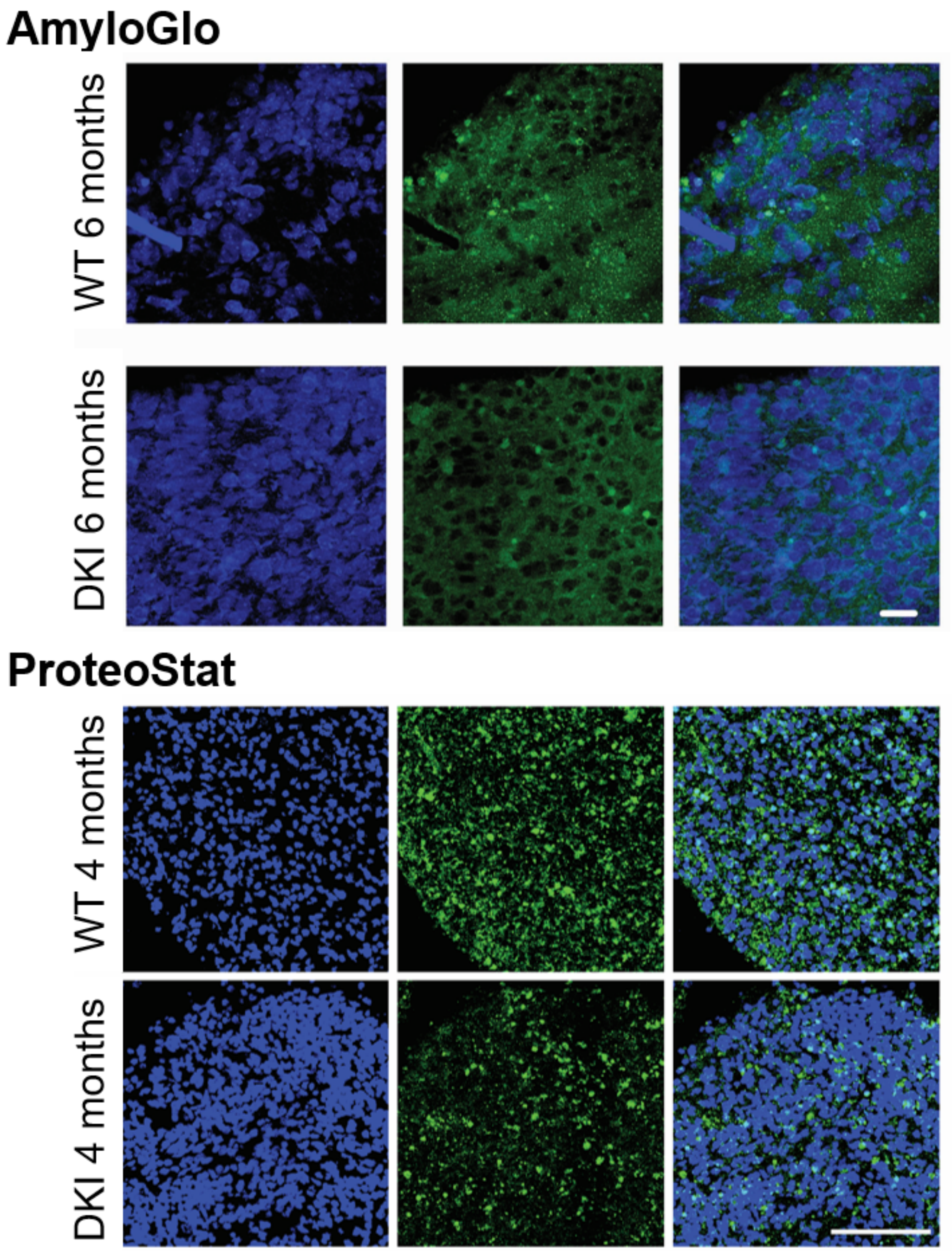


**Supplemental Figure 3: Representative Images of AmyloGlo and ProteoStat staining of cerebrocortical organoid sections.** Both dyes become more fluorescent when incorporated into amyloid structures. In both the WT and DKI cerebrocortical organoids we see staining of amyloid structures. Hoechst (nuclei, blue), AmyloGlo and ProteStat (green). AmyloGlo scale bar is 10 µm. ProteoStat scale bar is 100µm.

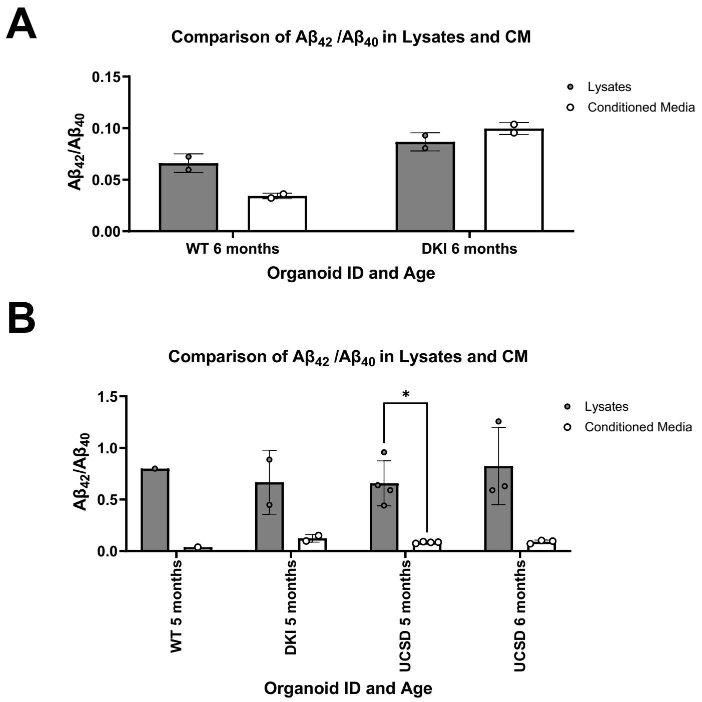


**Supplemental Figure 4. Comparison of $A\beta_{42}/A\beta_{40}$ ratios in matched organoids and conditioned media.** A) $A\beta_{42}/A\beta_{40}$ in WT conditioned media was greater than $A\beta_{42}/A\beta_{40}$ in organoid lysates of matched cultures at 6 months, while $A\beta_{42}/A\beta_{40}$ in DKI conditioned media was greater than $A\beta_{42}/A\beta_{40}$ in organoid lysates of matched cultures at 6 months. B) in a follow-up experiment, the $A\beta_{42}/A\beta_{40}$ measured in all lysate samples were elevated 10x from the expected range. Suspected result of experimental error, conclusions were not drawn from these data.

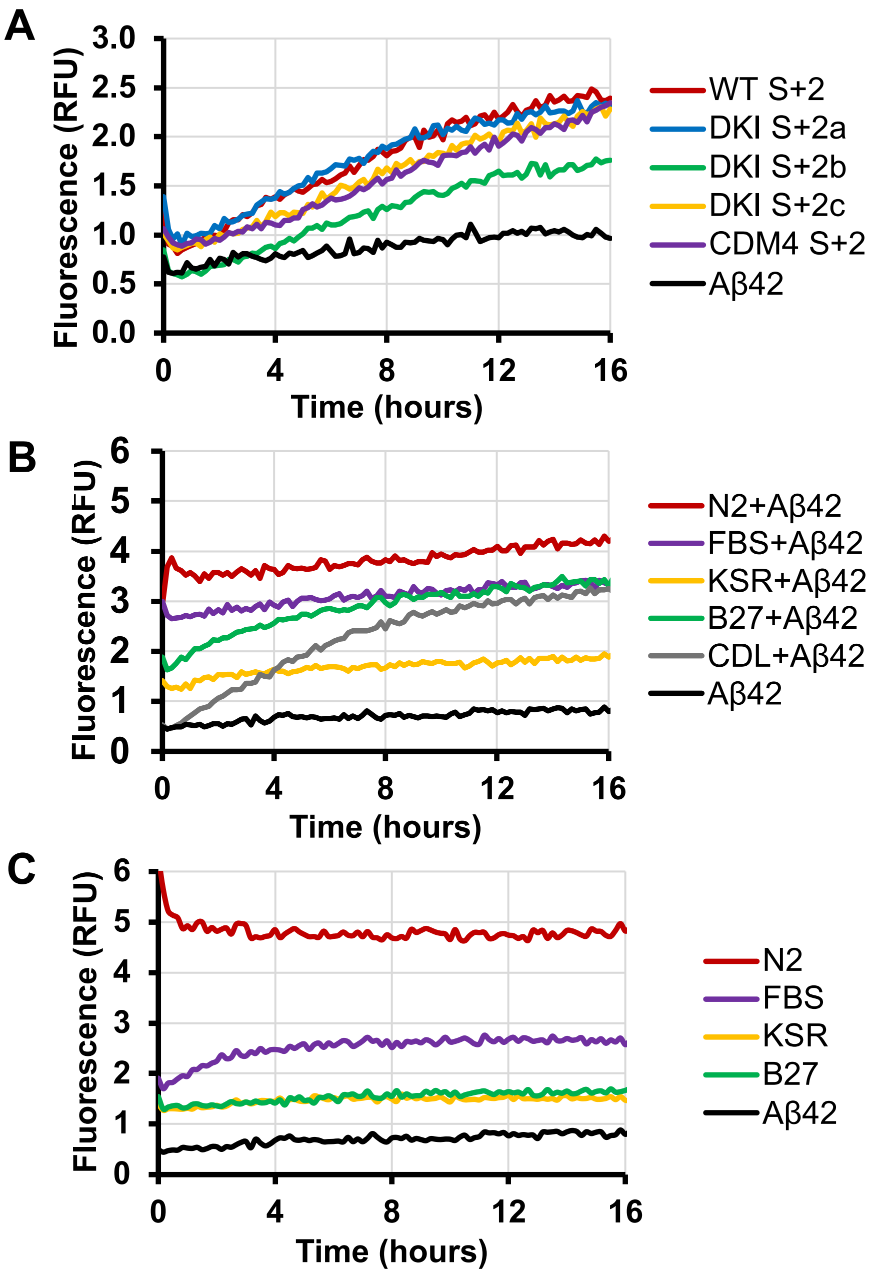


**Supplemental Figure 5: Amyloid-beta seed amplification assays.** A) Normalized Proteostat fluorescence curves showing similar seeding in matched fraction depth and volume CDM4 (processed media) versus DKI processed conditioned media. B) Proteostat fluorescence of common neuronal media components with amyloid beta peptide (Aβ42) over time. The increasing amounts of fluorescence show that all tested components initiate Aβ42 oligomerization to amyloid structures overtime. C) Proteostat fluorescence of common neuronal media components alone over time showing that media components alone interact with Proteostat amyloid dye. The increasing fluorescence of FBS indicates something in the FBS in making amyloid structures over time.
